# Striatal dopamine contributions to skilled motor learning

**DOI:** 10.1101/2024.02.06.579240

**Authors:** Chris D. Phillips, Courtney C. Myers, Daniel K. Leventhal, Christian R. Burgess

**Affiliations:** Michigan Neuroscience Institute, University of Michigan, Ann Arbor, MI, USA, 48109; Department of Molecular and Integrative Physiology, University of Michigan, Ann Arbor, MI, USA, 48109; Department of Neuroscience, University of Texas at Dallas, Richardson, TX, USA, 75080; Neuroscience Graduate Program, University of Michigan, Ann Arbor, MI, USA, 48109; Department of Neurology, University of Michigan, Ann Arbor, MI, USA, 48109; Department of Biomedical Engineering, University of Michigan, Ann Arbor, MI, USA, 48109; Parkinson Disease Foundation Research Center of Excellence, University of Michigan, Ann Arbor, MI, USA, 48109; Department of Neurology, VA Ann Arbor Health System, Ann Arbor, MI, USA, 48109

## Abstract

Coordinated multi-joint limb and digit movements - “manual dexterity” - underlie both specialized skills (e.g., playing the piano) and more mundane tasks (e.g., tying shoelaces). Impairments in dexterous skill cause significant disability, as occurs with motor cortical injury, Parkinson’s Disease, and a range of other pathologies. Clinical observations, as well as basic investigations, suggest that cortico-striatal circuits play a critical role in learning and performing dexterous skills. Furthermore, dopaminergic signaling in these regions is implicated in synaptic plasticity and motor learning. Nonetheless, the role of striatal dopamine signaling in skilled motor learning remains poorly understood. Here, we use fiber photometry paired with a genetically encoded dopamine sensor to investigate striatal dopamine release as mice learn and perform a skilled reaching task. Dopamine rapidly increases during a skilled reach and peaks near pellet consumption. In dorsolateral striatum, dopamine dynamics are faster than in dorsomedial and ventral striatum. Across training, as reaching performance improves, dopamine signaling shifts from pellet consumption to cues that predict pellet availability, particularly in medial and ventral areas of striatum. Furthermore, performance prediction errors are present across the striatum, with reduced dopamine release after an unsuccessful reach. These findings show that dopamine dynamics during skilled motor behaviors change with learning and are differentially regulated across striatal subregions.

## Introduction

Manual dexterity is a critical evolutionary adaptation for human survival. From the perspective of modern humans, dexterity is essential for everyday tasks like fastening buttons and tying shoelaces; however, stroke, multiple sclerosis, spinal cord injury, and several movement disorders impair dexterity (Alberts and Wolf, 2009; Benedict et al., 2011; Kamm et al., 2012; Nardone et al., 2013; Térémetz et al., 2015). In particular, people with Parkinson Disease (PD) experience “coordinative” deficits that significantly impair quality of life (Vanbellingen et al., 2018). Unlike bradykinesia and hypometria, coordinative deficits respond only partially and inconsistently to levodopa (Melvin et al., 2005; Negrotti et al., 2005; Doan et al., 2008; Lukos et al., 2013; Lee et al., 2018; Fasano et al., 2022). This suggests that aspects of dopamine signaling not restored by levodopa influence dexterous skill.

Rodent single pellet reaching (“skilled reaching”) is a commonly-used model of human dexterity in which rodents reach with a single forepaw towards a sugar pellet, grasp it, and retrieve it. Skilled reaching requires precise multi-joint coordination, depends on motor cortex for optimal performance (Whishaw et al., 1986; Guo et al., 2015), is homologous to human reaching (Sacrey et al., 2009), and recapitulates many features of human neurological disorders (Klein et al., 2012). Given the importance of motor cortex to human motor control, skilled reaching is highly likely to have translational relevance to human dexterous skill in health and disease.

Rodent skilled reaching is sensitive to dopamine manipulations. Rats with large 6-OHDA lesions make hypometric reaches similar to humans with PD (Whishaw et al., 1986, 2002). Coordination between proximal forelimb and distal digit movements are altered by acute dopamine manipulations specifically during reaches. Repeated optogenetic nigral stimulation or inhibition gradually accelerated or delayed transitions between reach submovements, respectively (Bova et al., 2020). These artificial and perhaps nonphysiologic manipulations demonstrate that altered nigrostriatal signaling influences not only reward processing, but also forelimb-digit coordination. It remains unknown, however, how striatal dopamine dynamics normally contribute to the acquisition and maintenance of fine motor skills.

We used fiber photometry recordings of the dopamine-sensitive biosensor dLight to determine striatal dopamine dynamics as mice learned and executed a head-fixed single-pellet reaching task. Dynamic dopamine signals emerged throughout the striatum, with a sharp increase at reach onset that peaked after the time the mouse made contact with the pellet. There were subtle but clear differences across striatal subregions, with broader peaks in ventral striatum and narrower peaks in dorsolateral striatum. Furthermore, ventral dopamine increased more than dorsal dopamine for cues predicting target availability. Somewhat surprisingly, these signals were nearly identical for the striatum ipsi- and contra-lateral to the reaching paw. Finally, there was a brief post-reach time window in which dopamine signals reflected successful vs failed grasping. These data show that striatal dopamine is modulated in specific ways across striatal subregions during skilled forelimb movements, and suggest a mechanism by which disrupted dopamine signaling could affect multi-joint coordination independently of movement “vigor.”

## Methods

### Mice

All animal care and experimental procedures were approved by the National Institute of Health and University of Michigan Institutional Animal Care and Use Committee. C57/B6 mice (https://www.jax.org/strain/000664, RRID:IMSR_JAX:000664) of both sexes (n = 24, aged 10-15 weeks at surgery) were housed at 22–24 °C with a 12 h light:12 h dark cycle with standard mouse chow and water provided *ad libitum*, unless otherwise stated.

### Surgery

Mice were anesthetized with isoflurane (2-3%) and placed into a stereotaxic apparatus (KOPF Model 963). For postoperative care, mice were injected intraperitoneally with meloxicam (5 mg/kg). After exposing the skull via small incision, a small hole was drilled through the skull for injection. A pulled-glass pipette with ∼20 μm tip diameter was inserted into the brain and virus was injected by an air pressure system. The pipette was kept in place for 5 min after injection before withdrawal. For *in vivo* photometry experiments, pAAV-CAG-dLight1.1 a gift from Lin Tian (titer 1.3 × 10^12^ genome copies per ml; Addgene viral prep # 111067-AAV1; http://n2t.net/addgene:111067; RRID:Addgene_111067) or pAAV-hSyn-EGFP a gift from Bryan Roth (titer ≥ 7×10¹² genome copies per ml; Addgene viral prep # 50465-AAV1; http://n2t.net/addgene:50465; RRID:Addgene_50465) were injected into the dorsolateral (100-200 nl, A/P: 0.9 mm, D/V: -2.9 mm, M/L: 2.1 mm from bregma), dorsomedial (100-200 nl, A/P: 0.9 mm, D/V: -2.9 mm, M/L: 1.1 mm from bregma), or ventral striatum (100-200 nl, A/P: 0.9 mm, D/V: -4.4 mm, M/L: 1.1 mm from bregma).

Optic fiber implantations were performed during the same surgery as viral injection. Metal ferrule optical fibers (400 μm diameter core, NA = 0.50; ThorLabs) were implanted. In a subset of mice, injections and fibers were implanted bilaterally (n = 9). Fibers were fixed to the skull using dental acrylic. Mice were given a minimum of 3 weeks recovery and 1 week acclimation before being used in any experiments. After the completion of the experiments, mice were perfused and the approximate locations of fiber tips were identified based on the coordinates of Franklin and Paxinos (Paxinos and Franklin, 2012).

### Experimental Design and Statistical Analysis

#### Skilled reaching training

Mice were first habituated by handling them for 5-10 minutes for 5 days. Mice were then food restricted to ∼85% of their *ad lib* body weight. Mice were acclimated to pellets (20 mg dustless precision pellets; BioServe) used in subsequent experiments. Mice were also habituated to head fixation by increasing the duration of head fixation each day across 5 days (10 minutes on day 1, 20 minutes on day 2, etc.). Paw preference was identified by holding food pellets in front of the head fixed mouse with forceps, allowing them to reach out and grab the pellet. Each mouse was then head fixed in the reaching apparatus with the pellet platform (Figure 1) close enough to their mouth that they could retrieve the pellet with their tongue or by scooping it into their mouth with their paw. Once mice were comfortable with this, the mouse was moved further from the pellet platform such that they had to reach out and grab the pellet. In a subset of mice that were reluctant to reach, we allowed a small number (<10) of reaches for pellets in a freely moving context before transitioning them again to the head fixed set up.

**Figure 1.**
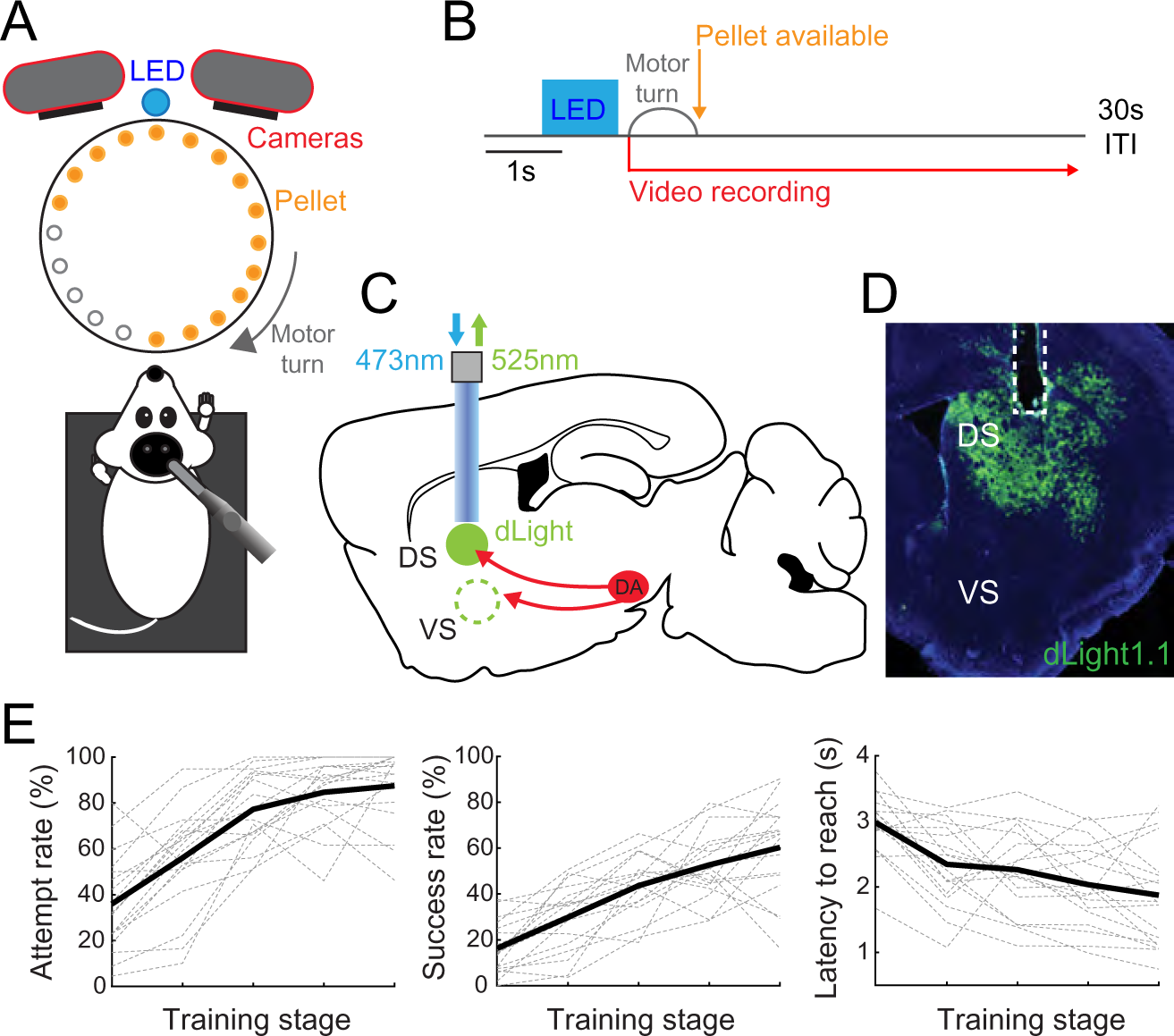
Mice learn and perform head-fixed skilled reach-to-grasp movements. A-B. Head-fixed mice perform a reach-to-grasp movement to obtain a food reward. An LED turns on for 1 s at trial onset, then a motor turns to place a food pellet within reach of the mouse. High-speed cameras acquire frames at 240 frames per second to track reach timing. A 30 s inter-trial interval follows each trial. C-D. To record dopamine dynamics in the striatum, dLight1.1 was expressed in striatal neurons and an optic fiber was placed to perform fiber photometry. E. Across training mice improved the proportion of trials where they attempted a reach (Left) and successfully reached (Middle). They also reduced their latency from LED offset to reaching onset (Right).

Trial start was signified by a LED turning on for 1s, followed by a stepper motor turning to place a pellet in front of the mouse (1.2 s turn duration; controlled via custom Arduino code). When the stepper motor turns, 2 high speed cameras (NaturalPoint, Inc., Optitrack PrimeX 13) are triggered to begin video acquisition at 240 f.p.s. for 6 s. After a 30 s inter-trial interval the next trial starts regardless of whether the mouse has reached for the pellet. 20 trials are performed per run and up to 4 runs occurred per day. Videos were manually scored to identify the trial outcome: ‘Non-attempt’ - where the mouse did not attempt a reach, ‘Attempt’ - where the mouse attempted a reach but did not successfully grasp the pellet, ‘Miss’ - where the mouse successfully grasped the pellet but did not successfully consume the pellet, and ‘Hit’-where the mouse successfully consumed the pellet. The mouse’s initiation of reaching, contact with the pellet, and consumption of pellet were manually scored from video frames. In a subset of experiments (n = 3), mice that were successfully reaching with their preferred paw had a physical blockade put in place to stop them from reaching, forcing them instead to reach-to-grasp with their other paw.

#### dLight photometry

A fiber optic patch cable (1 m long, 400 μm diameter; Doric Lenses) was firmly attached to the implanted fiber optic cannula with a zirconia sleeve (Doric Lenses). LEDs (473 nm; Plexon) were set such that a light intensity of <0.1 mW entered the brain; light intensity was kept constant across sessions for each mouse. Emission light was passed through a filter cube (Doric Lenses) before being focused onto a sensitive photodetector (2151, Newport). The signal was digitized with a National Instruments data acquisition card and collected using a custom MATLAB script. Fluorescent traces were bleaching corrected by subtracting a double exponential fit then adding back the mean of the trace prior to calculating ΔF/F. ΔF/F = ((F – F_0_)/F_0_; where F_0_ was calculated as the 10th percentile of the entire fluorescence trace was calculated for each session. These traces were z-scored to facilitate comparisons across days and mice. To establish differences in fluorescent time courses across groups we downsampled the data from 1000 Hz to 20 Hz. To quantify discrete changes locked to a given behavior in bar plots we established a baseline as the 1 s prior to the trial onset cue and compared it to the period immediately after trial onset (2 s), reach onset (1 s), pellet contact (1 s) or consumption (1 s).

To investigate dopamine responses in dorsal and ventral striatum during running and in response to liquid rewards, a subset of mice were head-fixed on a running wheel and given access to a lickspout with 10% sucrose. Wheel running was quantified by IR-beam breaks and licking behavior was quantified by a capacitive lickspout. Fluorescent signals were averaged at the onset of each running or licking bout then averaged across mice. We defined baseline as the 2 s prior to the behavior onset, running behavior as the 8 s after running onset, and licking behavior as the 2 s after licking onset.

#### Software accessibility

Analysis was performed using custom Matlab scripts. Basic analysis code is available via online repository (https://github.com/cburgess23/Mouse_dLight_skilledreaching.git).

#### Histology

Following the conclusion of all the experiments, mice were anesthetized with pentobarbital and transcardially perfused with phosphate-buffered saline (PBS) followed by 10% Formalin. Then, brains were post-fixed in 10% Formalin for 24h. Next, brains were transferred into 20% sucrose for ∼48 hours. Brains were then frozen and coronal sections were cut at 40 μm by a freezing microtome. Slices were then washed in PBS and incubated in a conjugated GFP rabbit polyclonal antibody as a primary antibody (1:5000 dilution, Novus, Littleton, CO; Catalog #: NB600-308AF488) at 4℃ overnight. The next day, slices were washed in PBS and mounted on glass slides using Vectashield - Antifade Mounting Medium with DAPI (Vector Laboratories, Burlingame, CA). Images were then captured by a fluorescence microscope (Olympus IX83) at 10x magnification.

#### Statistics

Statistical analyses were performed in Graphpad Prism or Matlab with an alpha level of 0.05. The data presented met the assumptions of the statistical test employed. Comparisons between baseline and response within cohorts were evaluated using paired t-tests, comparisons between average responses across cohorts were evaluated using a 2-way ANOVA, differences in time courses across cohorts were evaluated using 2-way ANOVAs on data downsampled by 50 from original acquisition rate (from 1000 Hz to 20 Hz).

Exclusion criteria for experimental animals were a) sickness or death during the testing period or b) if histological validation of the injection site demonstrated an absence of dLight or GFP expression. N-numbers represent the final number of healthy/validated animals.

## Results

### Striatal dopaminergic release is elevated during execution of a skilled reach

To identify dopamine release dynamics across skilled reaching, food-restricted mice were head-fixed in front of a pellet platform and trained to reach for the available pellets. Trial onset was signified by illumination of an LED for 1 s, then rotation of the platform to place a pellet in front of the mouse (Figure 1A and B). On each trial, the mouse may or may not reach for the pellet; therefore trial and reach counts are not necessarily the same. High-speed cameras captured 240 f.p.s. video to allow for scoring of mouse behavior. dLight1.1 was expressed in the striatum and an optic fiber was placed above the area of expression to allow for fiber photometry recordings of dopamine release (Figure 1C and D). Head-fixed mice (n=18) consistently performed a skilled reach-to-grasp movement. Over many trials (range: 140-440; mean: 290 trials/mouse) mice increased their attempt and success rates, culminating in a greater than 60% average success rate once well trained (Figure 1E; all trials split into five bins of equal trial numbers across learning for each mouse). Throughout training the latency to start a reach decreased (Figure 1E).

For mice with dLight1.1 expression and a fiber implanted in the striatum on the contralateral side to the reaching paw, we investigated striatal dopamine dynamics in response to trial onset (i.e., the LED), the start of the reach, contact with pellet, and start of consumption for successful hit trials (Figure 2A). There was a robust increase in dopamine-dependent fluorescence at trial onset; dopamine release increased at the start of the reach, peaking just prior to pellet consumption (Figure 2B; p < 0.05, 2-way ANOVA Sidak multiple comparisons test). Dopamine significantly increased at both trial onset (baseline: -0.09 +/- 0.02, response: 0.32 +/- 0.09; p < 0.001, 2-way ANOVA Sidak multiple comparisons test, Event: p = 0.005 F = 8.6, Group: p = 0.1 F = 2.7 Interaction: p = 0.006 F = 8.3) and at reach onset (baseline: -0.09 +/- 0.02, response: 1.43 +/- 0.14; p < 0.0001, 2-way ANOVA Sidak multiple comparisons test, Event: p < 0.0001 F = 45, Group: p < 0.0001 F = 26 Interaction: p < 0.0001 F = 35), when compared to the baseline period before the trial start (Figure 2C and D). No increase was observed in control mice expressing GFP instead of dLight (n = 6; Trial onset - baseline: 0.007 +/- 0.041, response: 0.007 +/- 0.039; p = 0.98; Reach - baseline: 0.007 +/- 0.041, response: 0.1 +/- 0.13; p = 0.57).

**Figure 2.**
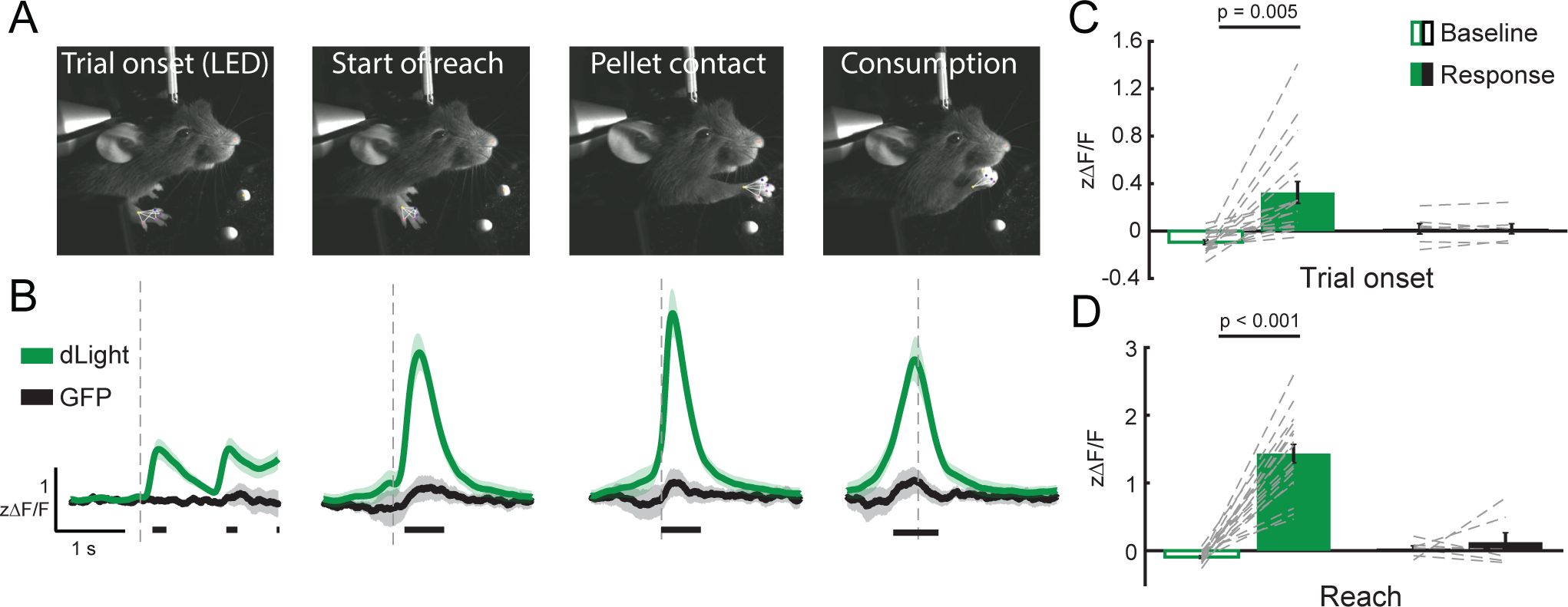
Striatal dopamine dynamics during skilled reaching. A. Using high-speed videography, task-specific events were scored, including the trial onset, the start of the mouse’s reach, the mouse grasping the pellet, and the onset of pellet consumption. B. Dopamine increased in response to the cues predicting trial onset and in response to reach onset, peaking around the time of pellet consumption. Lines under time courses denote a significant difference from GFP at that time, p<0.05 2-way ANOVA Sidak multiple comparisons test. C-D. dLight recordings demonstrated significant increases in response to trial onset and reach onset compared to baseline and compared to GFP control recordings.

We hypothesized that the reach-related dopamine response would be more prominent and sustained in the striatum contralateral to the reaching paw. In a subset of mice we recorded bilateral striatal dopamine release throughout behavioral training (n = 8). Dopamine levels increased in both the contralateral (Trial onset - baseline: -0.06 +/- 0.03, response: 0.27 +/- 0.10; p = 0.02, 2-way ANOVA Sidak multiple comparisons test, Event: p = 0.0005 F = 14, Group: p = 0.6 F = 0.26, Interaction: p = 0.88 F = 0.02; Reach - baseline: -0.06 +/- 0.03, response: 1.09 +/- 0.17; p < 0.01, 2-way ANOVA Sidak multiple comparisons test, Event: p < 0.0001 F = 66, Group: p = 0.36 F = 0.8, Interaction: p = 0.45 F = 0.55) and ipsilateral (Trial onset - baseline: -0.04 +/- 0.04, response: 0.20 +/- 0.10; p = 0.05; Reach - baseline: -0.04 +/- 0.04, response: 1.35 +/- 0.25; p = 0.001) striatum in response to trial onset and to the start of the reach (Figure 3A-B). There was no significant difference between contralateral and ipsilateral dopamine levels during either trial onset or reach onset (Trial onset: p = 0.88, F = 0.02; Reach: p = 0.36, F = 0.84). In a further subset of mice we blocked their preferred reaching paw after they had become proficient in reaching, forcing them to learn to reach with their non-preferred paw (n = 3; Figure 3C-D). These mice achieved similar Hit rates with their less preferred paw. During both normal and paw-blocked reaching, mice showed a significant increase in dopamine release at reach onset in both the contralateral and ipsilateral striatum relative to the reaching paw (normal reaching: p<0.01, 2-way ANOVA Sidak multiple comparisons test, Event: p < 0.0001 F = 95, Group: p = 0.64 F = 0.22, Interaction: p = 0.74 F = 0.11; paw-blocked reaching: p<0.01, 2-way ANOVA Sidak multiple comparisons test, Event: p < 0.0001 F = 92, Group: p = 0.94 F = 0.004, Interaction: p = 0.66 F = 0.02).

**Figure 3.**
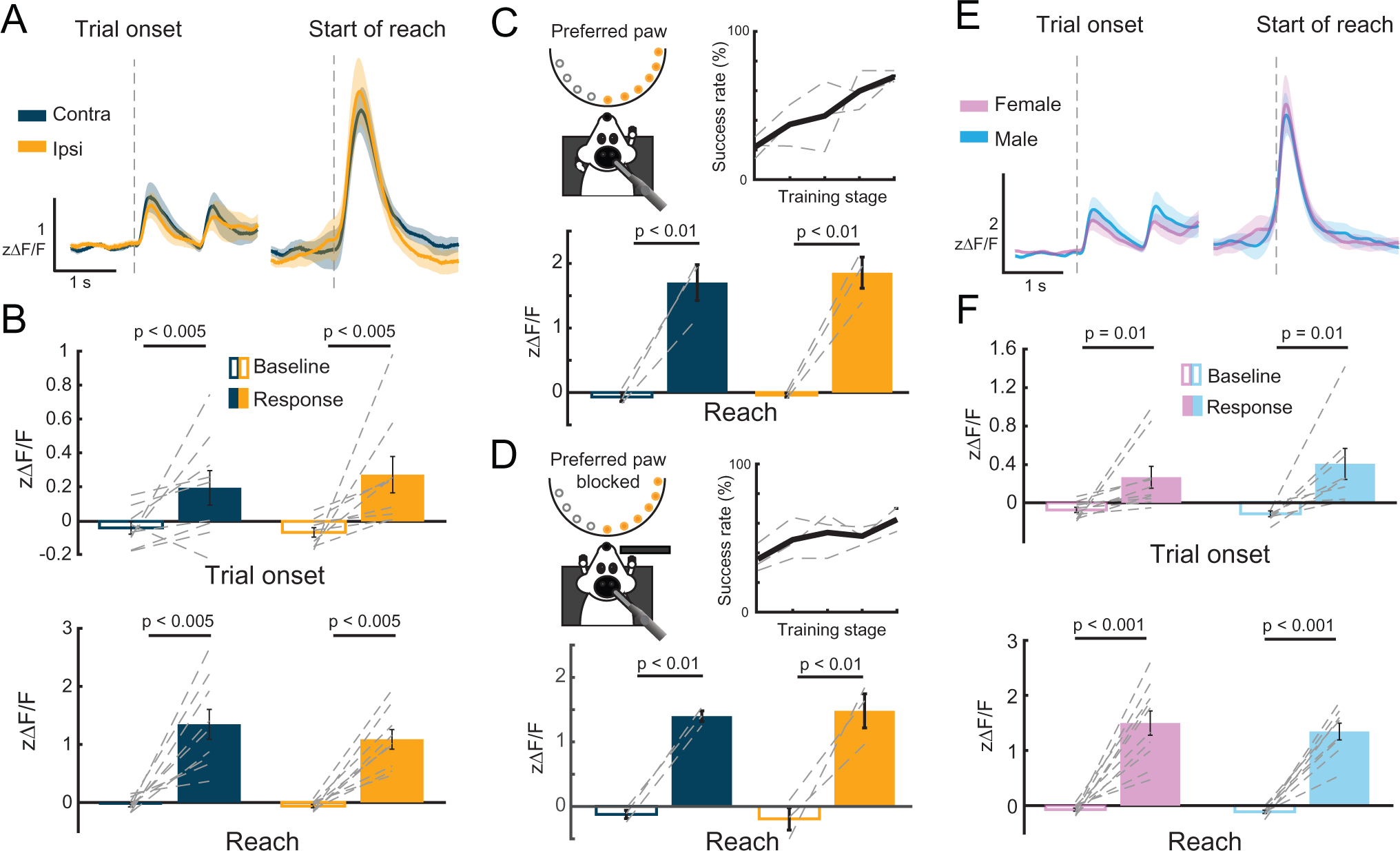
Striatal dopamine dynamics are similar in both contra- and ipsi-lateral striatum. A. Dopamine dynamics were similar in the contralateral striatum and ipsilateral striatum relative to the reaching paw. B. dLight recordings demonstrated significant increases in response to trial onset and reach onset compared to baseline in both the contralateral and ipsilateral striatum. C. Mice initially learned to reach with their preferred reaching paw and perform the task well. Dopamine significantly increased during reaching relative to baseline in both the contralateral and ipsilateral striatum relative to the reaching paw. D. Mice in which their preferred reaching paw was physically blocked could re-learn the task and perform well. Dopamine significantly increased during reaching relative to baseline in both the contralateral and ipsilateral striatum relative to the new, non-prefered, reaching paw. E. Dopamine increased in response to the cues predicting trial onset and in response to reach onset similarly in both male and female mice. F. Recordings showed no significant difference between male and female mice in response to trial onset and reach onset.

Dopamine levels increased in both the female (n = 10; Trial onset - baseline: -0.07 +/- 0.03, response: 0.26 +/- 0.11; p = 0.03, 2-way ANOVA Sidak multiple comparisons test, Event: p < 0.0001 F = 24, Group: p = 0.6 F = 0.26, Interaction: p = 0.33 F = 0.94) and male (n = 8; Trial onset - baseline: -0.11 +/- 0.03, response: 0.40 +/- 0.16; p = 0.05) mice in response to trial onset (Figure 3E-F). Similarly, both female (Reach - baseline: - 0.07 +/- 0.03, response: 1.49 +/- 0.22; p < 0.0001, 2-way ANOVA Sidak multiple comparisons test, Event: p < 0.0001 F = 20, Group: p = 0.5 F = 0.08, Interaction: p = 0.72 F = 0.02) and male (Reach - baseline: -0.11 +/- 0.03, response: 1.35 +/- 0.15; p < 0.0001) mice had significantly elevated dopamine responses at reach onset. There was no significant difference between male and female mice during either trial or reach onset (Trial onset: p = 0.61, F = 0.26; Reach: p = 0.51, F = 0.45).

Previous recordings of dopamine activity suggest that dopamine dynamics may vary across striatal subregions, particularly along the dorsal-ventral axis (Howe and Dombeck, 2016; Parker et al., 2016). We confirmed this by quantifying dopamine release in response to spontaneous bouts of running or licking for a sucrose reward. While there were some dynamics to both events in dorsal striatum (n = 6; Running - baseline: 0.21 +/- 0.03, response: 0.32 +/- 0.04, p = 0.05, paired t-test; Licking - baseline: 0.35 +/- 0.07, response: 0.45 +/- 0.11, p = 0.41, paired t-test) and ventral striatum (n = 4; Running - baseline: 0.26 +/- 0.06, response: 0.36 +/- 0.02, p = 0.2, paired t-test; Licking - baseline: 0.02 +/- 0.07, response: 0.89 +/- 0.12, p < 0.005, paired t-test), there was a bias towards reward type responses in ventral striatum and motor responses in dorsal striatum (Figure 4A-B). No dynamics were seen in response to running or licking in GFP controls (n = 5; Figure 4C).

**Figure 4.**
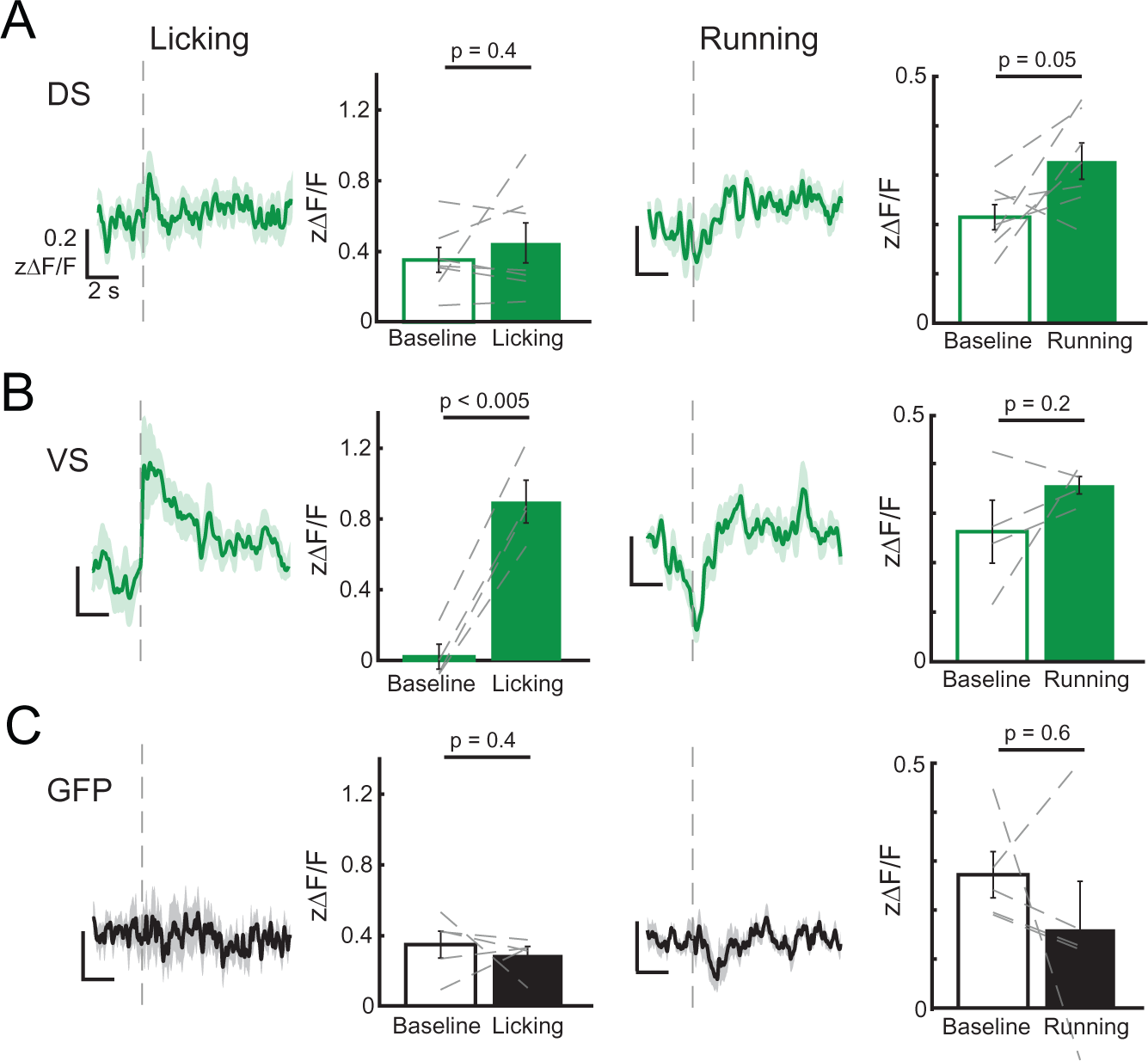
Dopamine release differs across the dorsal-ventral axis of the striatum. A-B. Dopamine increases in response to locomotion and liquid rewards, with recordings in the dorsal striatum responding more strongly to running, and ventral recordings responding more strongly to rewards. C. GFP controls showed no response to either behavior.

### Striatal subregion specific dopamine dynamics during skilled reaching behavior

We separated our recordings into mice with fibers in the dorsolateral striatum (DLS; n = 4), dorsomedial striatum (DMS; n = 6), and ventral striatum (VS; n = 8) on the hemisphere contralateral to the reaching paw (Figure 5A). We investigated dopamine responses in each area during trials that were successful (Hit trials), unsuccessful (Miss trials) in addition to trials where the mouse attempted to reach but failed to grasp the pellet (Attempt trials), and where the mouse did not attempt a reach (Non-attempt trials). In the DLS, dopamine was elevated slightly during trial onset cues and sharply at reach onset, with fast onset and offset dynamics (Figure 5B). There was a significant increase relative to baseline during both the trial onset response (baseline: -0.07 +/- 0.04, response: 0.11 +/- 0.03, p = 0.3, paired t-test) and the reach response (baseline: -0.07 +/- 0.04, response: 1.79 +/- 0.28, p < 0.01, paired t-test; Figure 5C-D). When comparing Hit and Miss trials there was evidence of performance prediction error, with reduced dopamine in the Miss trials at the time when pellet consumption would have happened in Hit trials (Hit: 0.62 +/- 0.25, Miss: -0.48 +/- 0.08, p = 0.01, paired t-test; Figure 5E and 5B pink shaded area).

**Figure 5.**
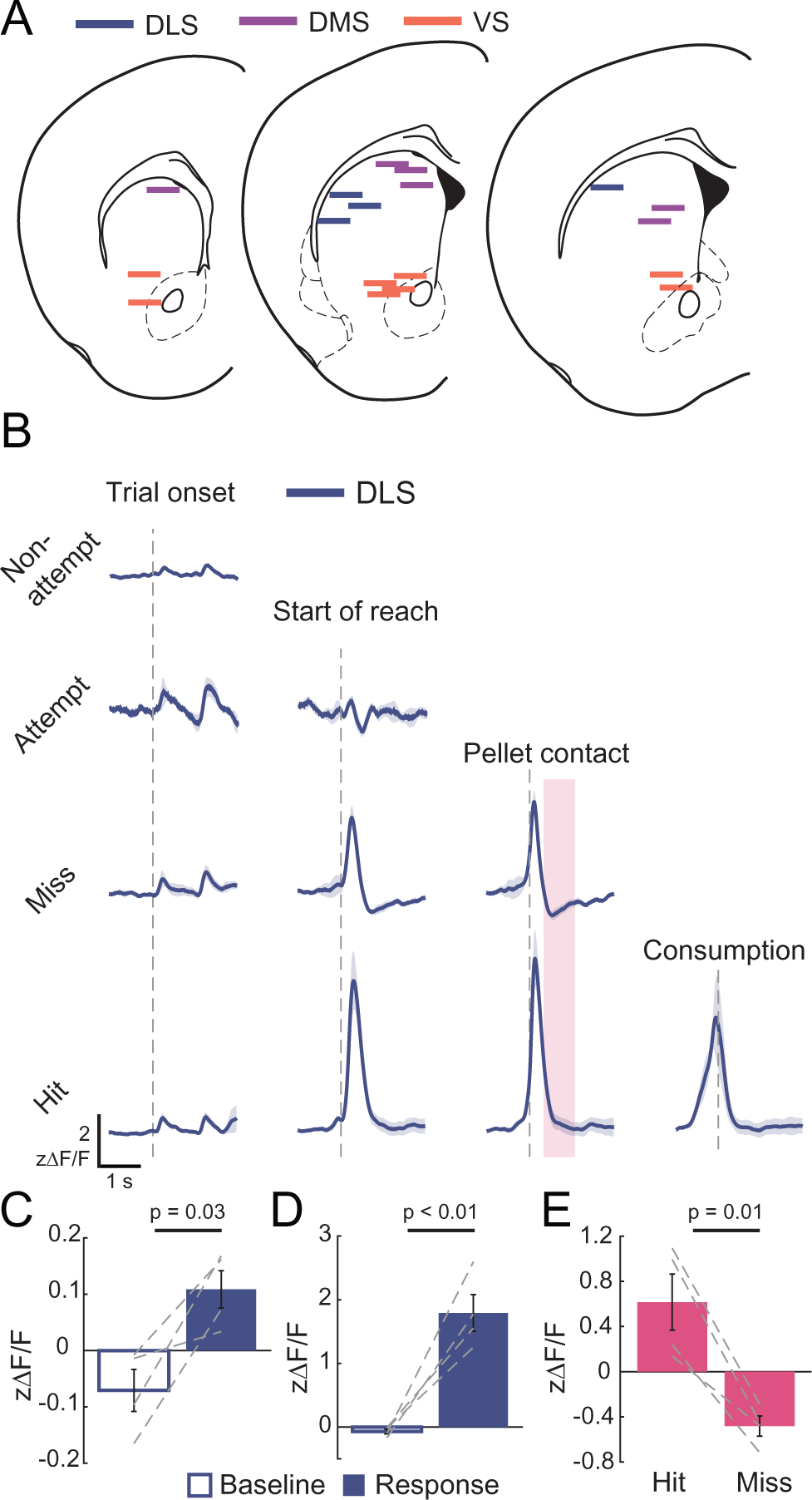
Dorsolateral striatal dopamine dynamics during skilled reaching. A. Fiber locations for photometry recordings across subregions of the striatum. B. Dopamine dynamics in DLS in response to Non-attempt trial, Attempt trial, Hit trial, and Miss trial outcomes. C-D. DLS dopamine increased in response to trial onset and reach onset when compared to baseline. E. Performance prediction error is evident in DLS when comparing late-reach dopamine dynamics in Hit vs. Miss trials (pink shaded area in A).

In the DMS, dopamine was elevated during trial onset cues and to the onset of a reach, with fast onset and slower offset dynamics (Figure 6A). There was a significant increase, relative to baseline, during the reach response (baseline: -0.06 +/- 0.03, response: 1.08 +/- 0.19, p = 0.002, paired t-test; Figure 6B-C). When comparing Hit and Miss trials there was clear evidence of performance prediction error, with a reduced dopamine response in the Miss trials at the time when pellet consumption would have happened in Hit trials (Hit: 0.78 +/- 0.19, Miss: -0.13 +/- 0.13, p = 0.01, paired t-test; Figure 6D and 6A pink shaded area).

**Figure 6.**
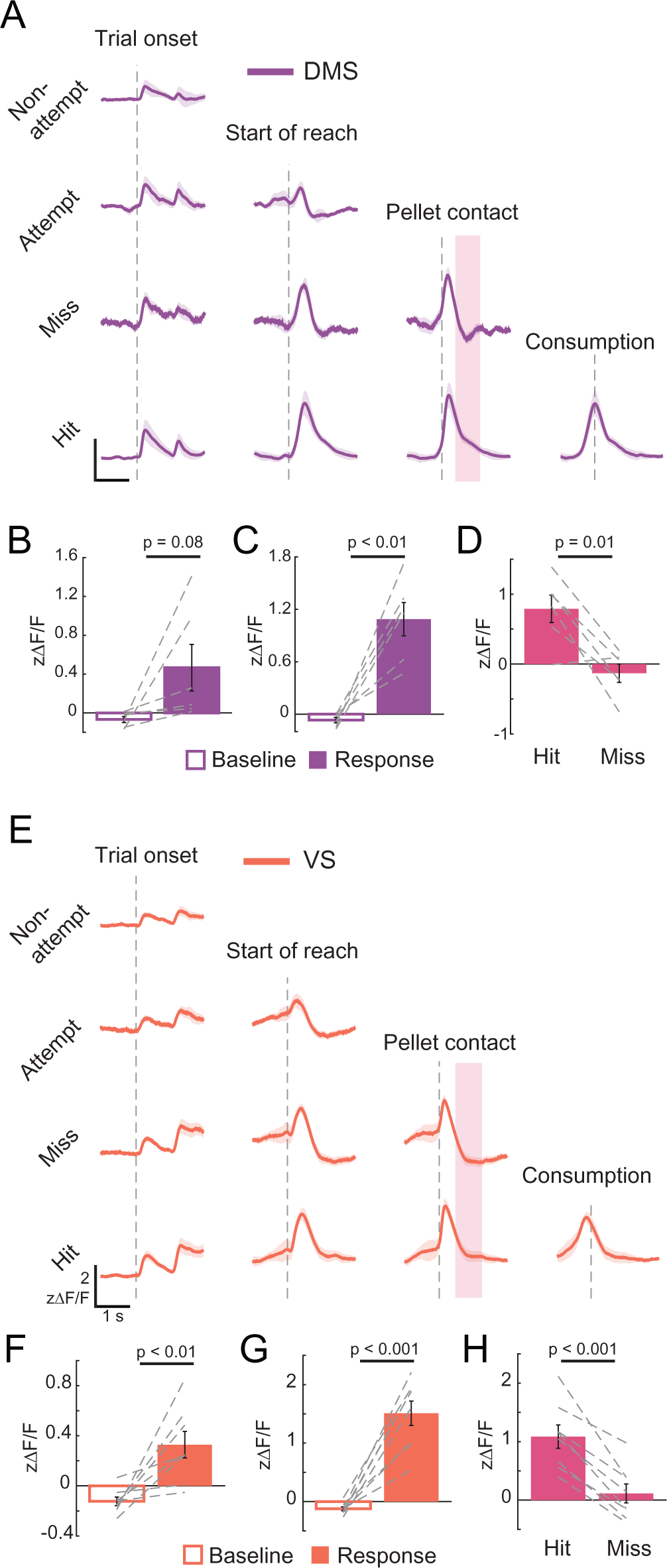
Dorsomedial and ventral striatal dopamine dynamics during skilled reaching. A. Dopamine dynamics in DMS in response to Non-attempt trial, Attempt trial, Hit trial, and Miss trial outcomes. B-C. DMS dopamine did not increase significantly in response to trial onset, showing variability across recordings, but did increase at reach onset when compared to baseline. D. Performance prediction error is evident in DMS when comparing late-reach dopamine dynamics in Hit vs. Miss trials (pink shaded area in E). E-H. Same as A-D but for ventral striatal dopamine release.

In the VS, dopamine was elevated during trial onset cues and to the onset of a reach, with slower onset and offset dynamics (Figure 6E). There was a significant increase relative to baseline during both trial onset (baseline: -0.12 +/- 0.03, response: 0.32 +/- 0.10, p = 0.007, paired t-test;) and the reach response (baseline: - 0.12 +/- 0.03, response: 1.51 +/- 0.20, p < 0.001, paired t-test; Figure 6F-G). When comparing Hit and Miss trials there was clear performance prediction error, with a reduced dopamine response in the Miss trials at the time where pellet consumption would have happened in Hit trials (Hit: 1.08 +/- 0.2, Miss: 0.11 +/- 0.16, p < 0.001, paired t-test; Figure 6H and 6E pink shaded area).

When comparing dopamine release during execution of skilled reaching across DLS, DMS, and VS there were different release dynamics across areas. VS dopamine was increased during the trial onset cues, specifically in response to the stepper motor turning, when compared to both DLS and DMS (Figure 7; 2-way ANOVA Sidak multiple comparisons test, p<0.05). VS dopamine had slower dynamics, both rising and falling more gradually than dorsal recordings. DLS dopamine peaked at a higher level than VS and DMS, and had generally faster dynamics (Figure 7; 2-way ANOVA Sidak multiple comparisons test, p<0.05).

**Figure 7.**
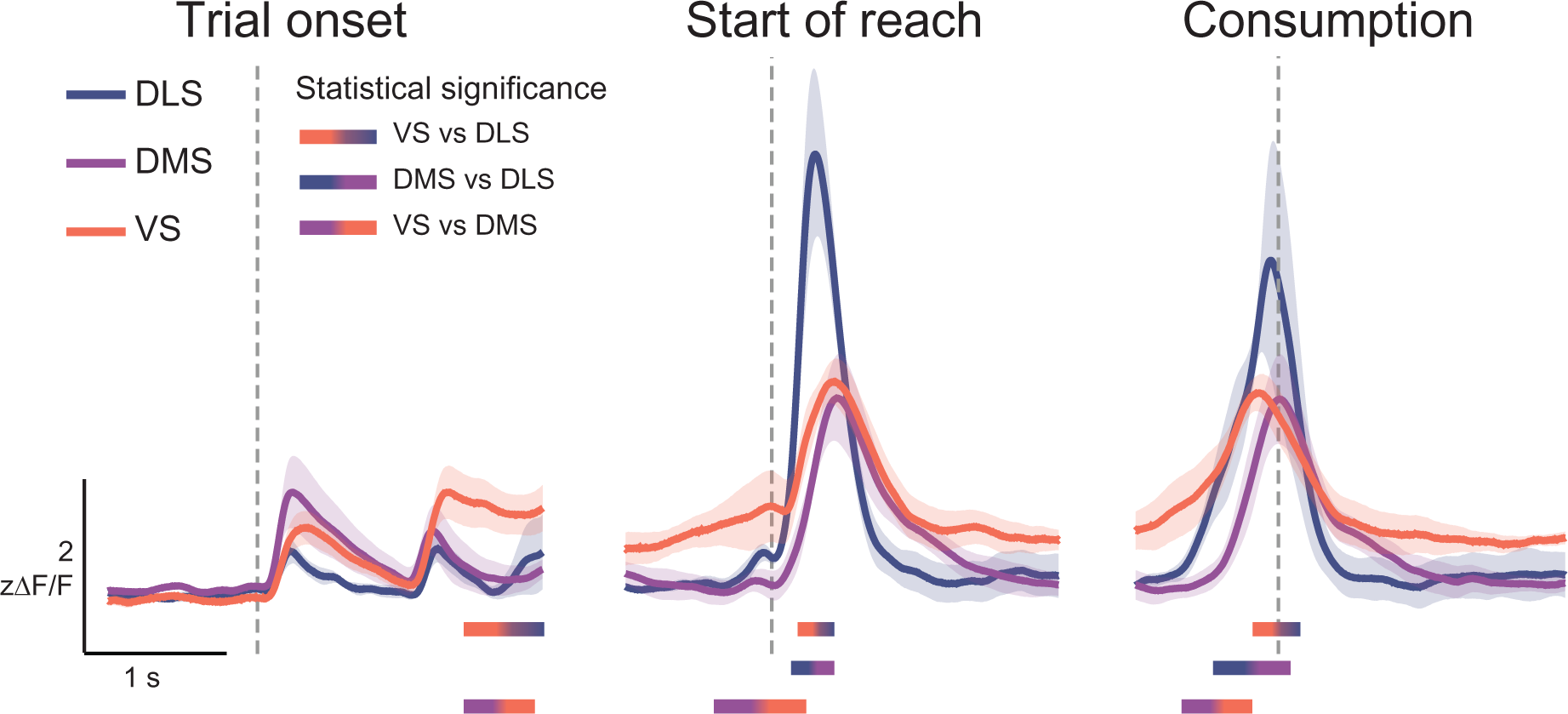
Differences in dopamine dynamics across striatal areas. Comparison of dopamine dynamics in different subregions of the striatum. Lines under time courses denote a significant difference at that time, p<0.05 2-way ANOVA Sidak multiple comparisons test.

### Dopamine dynamics change across motor learning

We next investigated how dopamine dynamics change across learning of skilled reaching behavior in different areas of the striatum. For each mouse the first twenty Hit trials (Early learning) were compared to the last twenty Hit trials (Late learning). In the DLS, peak reach dopamine response, half-peak width of the reach response, and the timing of the peak dopamine response relative to the start of the reach were all unchanged (p > 0.4, paired t-tests; Figure 8A). Example heatmaps from a representative mouse show that dopamine dynamics were largely similar across learning (Figure 8B). The response to trial onset increased slightly (Figure 8C; Early: -0.05 +/- 0.9, Late: 0.26 +/- 0.04, p = 0.04; paired t-test), but reach response and consumption response were similar across learning, resulting in only a subtle shift in dopamine release from the reach to the trial onset cues (Figure 8D).

**Figure 8.**
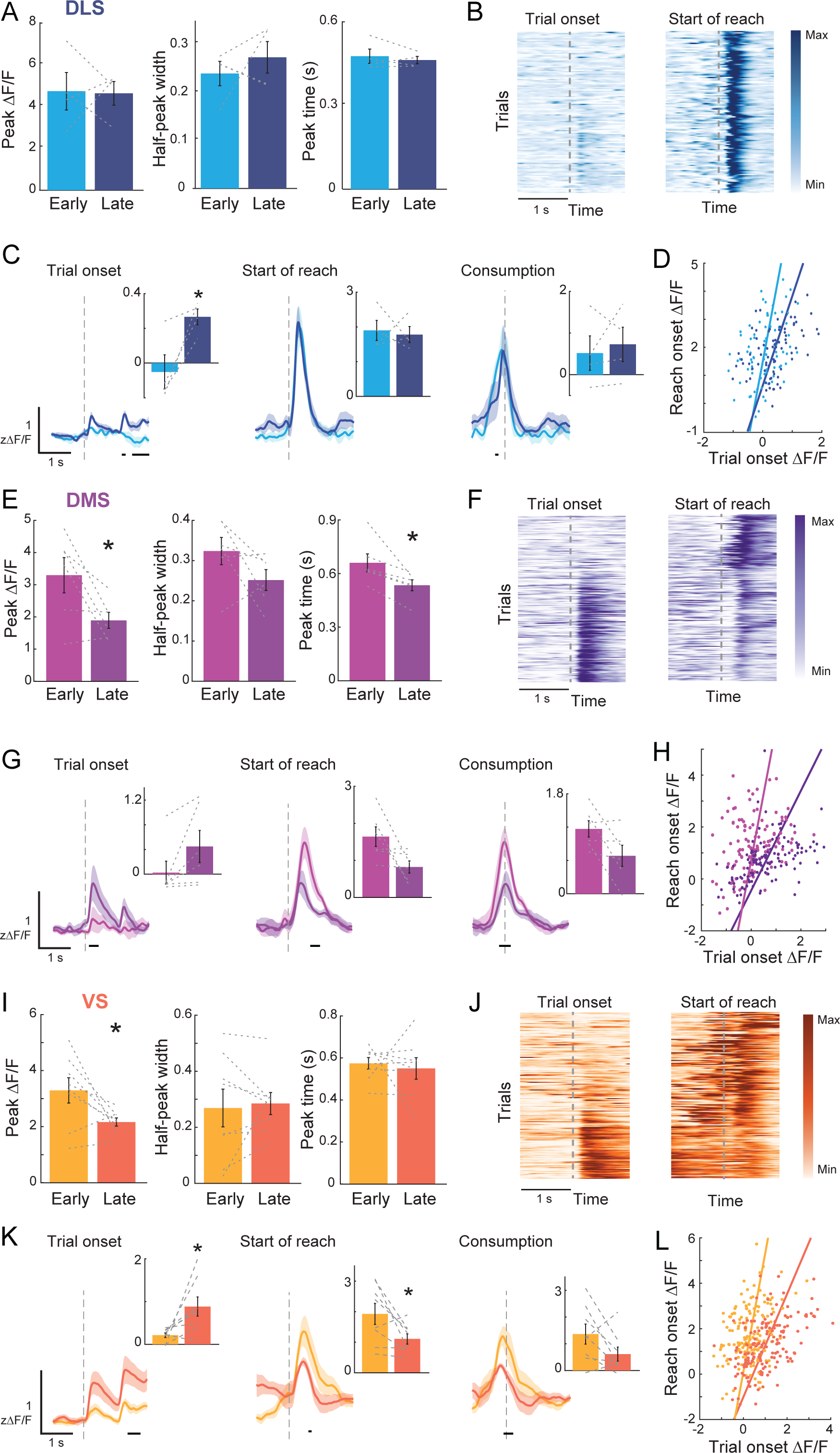
Dopamine dynamics change across learning of a reach-to-grasp movement. A. Peak dopamine reach response, half-peak width of the dopamine reach response, and the timing of the peak dopamine reach response across learning in DLS. B. Example heatmap from Hit trials across learning in one mouse with a DLS recording. C. Differences in dopamine dynamics across learning in DLS. D. Relationship between peak ΔF/F at trial and reach onset for early and late reaches. Dopamine responses tend to shift from reach onset to trial onset with training. E-H. Same as A-D for DMS. I-J. Same as A-D for VS. * denotes a significant difference, p<0.05 paired t-test. Lines under time courses denote a significant difference at that time, p<0.05 2-way ANOVA Sidak multiple comparisons test.

In the DMS, peak dopamine reach response was decreased in late vs. early learning (Figure 8E; early: 3.2 +/- 0.54, late: 1.9 +/- 0.25, p = 0.04; paired t-test), as was the timing of the peak dopamine response (early: 0.66 +/- 0.05, late: 0.53 +/- 0.03, p = 0.02; paired t-test); this shift in timing may reflect increased speed of successful reaches, as both early and late learning have a peak dopamine response at the time of consumption. Dopamine dynamics showed subtle differences between early and late learning. Example heatmaps from a representative mouse show that dopamine dynamics change across learning (Figure 8F).

Specifically, brief periods around LED onset showed an increased response during late learning and pellet consumption showed an increased response during early learning (Figure 8G), these changes were not significant when averaging over larger time periods (p> 0.05, paired t-tests; Figure 8G insets), but overall resulted in a shift in relative response from the reach to the trial onset cues (Figure 8H).

In the VS, peak dopamine reach response was decreased in late vs. early learning (Figure 8I; early: 3.3 +/- 0.45, late: 2.2 +/- 0.15, p = 0.04, paired t-test), while there was no change in half-peak width and the timing of the peak dopamine response (p>0.05, paired t-tests). Dopamine dynamics showed clear differences across learning in line with reward prediction errors. Example heatmaps from a representative mouse show that dopamine dynamics shift from the reach response to the cue predicting trial onset (Figure 8J). Trial onset cue responses increased during late learning and pellet consumption responses decreased during late learning (Figure 8K), resulting in a shift in relative response from the reach to the trial onset cues (Figure 8L). These changes were reflected when averaging over larger time periods for both trial onset (early: 0.21 +/- 0.05, late: 0.88 +/- 0.22, p = 0.03, paired t-test) and reach (early: 1.9 +/- 0.34, late: 1.1 +/- 0.17, p = 0.03, paired t-test; Figure 8K insets).

## Discussion

We found sharp bilateral striatal dopamine transients time-locked to reach-to-grasp movements in mice. Dopamine increased sharply at reach onset, with peaks locked most tightly to the pellet contact and consumption. These phasic dopamine signals varied in subtle but consistent ways across striatal subregions, with narrower peaks in DLS. These responses migrated to the cue signaling impending pellet availability, primarily in VS and DMS. Finally, dopamine levels remained relatively elevated in successful trials, but dropped below baseline with failed pellet retrieval. These results suggest that phasic striatal dopamine signaling plays an important role in learning and maintaining dexterous skill.

Striatal dopamine is generally considered to subserve two functions: regulating movement vigor and mediating reinforcement learning. Here, we use “vigor” to represent the amplitude, speed, or frequency of movement, which is intentionally broad to capture the range of deficits observed in dopamine-deficient states (Dudman and Krakauer, 2016). Over minutes to hours, tonic dopamine levels clearly influence one or more of these aspects of vigor (Mazzoni et al., 2007; Niv et al., 2007; Beeler et al., 2010b; Leventhal et al., 2014; Panigrahi et al., 2015). On shorter timescales corresponding to phasic signaling, however, the evidence is mixed. Several studies find that dopamine neurons increase their activity at or prior to movement onset, and/or dopamine neuron stimulation causes rapid movement initiation (Howe and Dombeck, 2016; Coddington and Dudman, 2018; da Silva et al., 2018; Saunders et al., 2018; Hunter et al., 2022). Others find that the primary dopamine neuron response to movement initiation is a firing pause (Dodson et al., 2016) or decrease in striatal dopamine or terminal activation (Tsutsui-Kimura et al., 2020; Markowitz et al., 2023). It therefore remains unclear if, or under what conditions, phasic dopamine release regulates instantaneous vigor.

If phasic peri-reach striatal dopamine signaling “invigorates” reaching, two things should be true. First, the absence of phasic dopamine signaling should prevent reaching, or at least significantly alter reach kinematics. Second, manipulating nigrostriatal dopamine should immediately alter reach kinematics. Regarding the first prediction, phasic peri-reach dopamine release in DMS and VS decreased with experience as reach accuracy improved. It is therefore unlikely that phasic dopamine release in these subregions influences instantaneous reach kinematics, though it is possible that DLS phasic dopamine release does. Previous optogenetic experiments also argue against an immediate effect of phasic dopamine signaling on forelimb kinematics. Repeated dopamine neuron stimulation or inhibition progressively altered forelimb kinematics across many reaches, but did not instantaneously affect reaching or grasping (Bova et al., 2020). This is consistent with a recent report that restoring tonic striatal dopamine levels permits non-dexterous forelimb movements in dopamine-depleted mice, but that phasic dopamine release is not required (Liu et al., 2022). Thus, it is unlikely that phasic striatal dopamine release regulates reach “vigor” on short timescales.

An alternative interpretation is that phasic dopamine facilitates corticostriatal plasticity to establish effective reach-to-grasp kinematics with repetition. By analogy with “reward prediction error” signals in Pavlovian or instrumental tasks (Schultz et al., 1997; Mohebi et al., 2019), striatal dopamine could signal a “performance prediction error” used to adapt reach strategy. This concept is consistent with dopamine neuron recordings in songbirds, in which changes in dopamine neuron activity reflect errors in song production (Duffy et al., 2022). Similarly, dopamine receptor blockade and dopamine replacement cause progressive rather than immediate changes in rotarod performance in mice (Beeler et al., 2010a, 2012). During spontaneous mouse behavior, striatal dopamine increases at the onset of repeatable behavioral modules, but this increase is not correlated with movement kinematics. Instead, it increases the probability that that behavior will be repeated (Markowitz et al., 2023; Tang et al., 2023). Our finding that post-reach dopamine levels correlate with reach outcome (“Miss” or “Hit”) is consistent with phasic dopamine as mediating skill acquisition across many trials.

Still, a pure “performance prediction error” signal is unlikely to explain the role of phasic striatal dopamine in motor learning for two reasons. First, there are more degrees of freedom in reach-to-grasp movements than instrumental tasks. Given a choice between discrete alternatives, a dopamine dip in the context of reinforcement learning has a clear interpretation: select a different option on subsequent trials. However, it is not obvious how a mouse should adjust its reach-to-grasp movement given a dopamine dip after a failed reach. The second reason is that optogenetic manipulation of nigrostriatal dopamine signaling during reaches causes predictable changes in forelimb kinematics (Bova et al., 2020). If phasic dopamine release reflects reach outcomes, reach kinematics should be more variable after peri-reach dopamine suppression and more consistent after peri-reach dopamine stimulation. Instead, dopamine neuron inhibition and stimulation had consistent but opposite effects on reach kinematics: stimulation shortened reaches and accelerated transitions between reach submovements, while inhibition lengthened reaches and delayed the transition from paw transport to grasping. These interventional experiments coupled with our new results indicate that phasic dopamine signaling in striatum plays a role in learning and adapting skilled movements. The precise meaning of these signals remains unclear, however.

Mechanistically, phasic dopamine release likely regulates corticostriatal plasticity at synapses active during reaches. D1 dopamine receptors along the striatal “direct” pathway have a lower binding affinity than indirect pathway D2 receptors. It is therefore suggested that phasic dopamine increases facilitate plasticity along the direct pathway, while phasic dopamine dips facilitate plasticity along the indirect pathway (Surmeier et al., 2007; Dreyer et al., 2010). More recent work, however, suggests that phasic changes in dopamine strongly affect both D1 and D2 receptors (Marcott et al., 2014; Yapo et al., 2017), with effects lasting much longer than the release event itself (Hunger et al., 2020). A dopamine pulse followed by a dip is predicted to quickly restore baseline D1 and D2 receptor occupancy, while a phasic increase followed by a return to baseline dopamine levels is predicted to cause a prolonged increase in D1 and D2 receptor occupancy. The phasic increase we observed at reach onset may prime dopamine receptors to facilitate corticostriatal plasticity when information regarding the outcome becomes available (Coddington and Dudman, 2019). Changes in corticostriatal synapses may underlie the increase in corticostriatal synchrony that occurs as skilled reaching is acquired (Lemke et al., 2019).

Reach-to-grasp movements recruit a complex network of cortical and subcortical structures. Motor cortex is important for accurate paw transport as well as fine digit movements necessary for grasping (Guo et al., 2015; Wang et al., 2017; Lemke et al., 2019; Sauerbrei et al., 2020; Park et al., 2022). Interestingly, motor cortical dopamine is necessary to learn, but not perform, single pellet reach-to-grasp (Hosp et al., 2011). Cortical control of paw transport depends on thalamic input (Sauerbrei et al., 2020), which in turn receives input from the basal ganglia and cerebellum. Basal ganglia, thalamus, and cerebellum are implicated in the control of paw transport, as opposed to fine digit control for grasping (Lemke et al., 2019; Guo et al., 2021; Lopez-Huerta et al., 2021; Calame et al., 2023). However, this may be because these studies tracked paw position, but not individual digits. The cerebellum seems to be important specifically for online adjustment of reach trajectories. Collectively, these studies suggest a model in which corticostriatal-thalamocortical circuits initiate a sequence of motor commands to transport the paw to the pellet and initiate a well-timed grasp. The phasic dopamine signals observed here presumably provide feedback to adjust kinematic parameters across reaches, while the cerebellum provides on-line adjustments during individual reaches. Further work will be required to validate this model and identify the specific aspects of reach-to-grasp movements modulated by dopamine, as well as the neural mechanisms by which they are implemented.

## Acknowledgments

We would like to thank Elizabeth Pappas for help training mice and Brandon Toth as well as other members of the Burgess lab for helpful discussion and feedback. This work was supported by a Brain Research Foundation Seed Grant and NIH R01DK129366 (CRB).

